# Development of a novel aging clock based on chromatin accessibility

**DOI:** 10.1101/2022.08.11.502778

**Authors:** Cheyenne Rechsteiner, Francesco Morandini, Kevin Perez, Viviane Praz, Guillermo López-García, Laura Hinte, Ferdinand von Meyenn, Alejandro Ocampo

**Affiliations:** Department of Biomedical Sciences, University of Lausanne, Switzerland; Department of Health Sciences and Technology, ETH Zurich, Switzerland; Department of Computer Sciences, University of Málaga, Spain

**Keywords:** aging, epigenetic clock, chromatin accessibility, ATAC sequencing

## Abstract

The establishment of aging clocks based on age-associated changes in DNA methylation has highlighted the strong link between epigenetic alterations and aging. However, the connection between DNA methylation changes at clock sites and their effect on cellular function remains unclear. We hypothesize that chromatin accessibility, a readout that integrates multiple epigenetic mechanisms, may connect epigenetic changes with downstream biological effects. To investigate this hypothesis, we generated chromatin accessibility profiles from peripheral blood mononuclear cells (PBMCs) of 157 human donors and construct a novel aging clock with a median absolute error on prediction of 5.69 years. Moreover, by comparing our chromatin accessibility data to matched transcriptomic profiles, we show that the genomic sites selected for the prediction of age based on chromatin accessibility undergo transcriptional changes during aging. This chromatin accessibility clock could therefore be used to investigate the direct effect of aged epigenetic states on cellular function.

## Background

Aging is a ubiquitous biological process that is characterized by a progressive loss of physiological integrity on multiple biological scales and increased vulnerability to disease and death [1]. Current global demographic trends toward an aged population highlight the importance of studying aging to understand its dynamics and mitigate its role as driver of diseases late in life.

The dysregulation of epigenetic networks is a crucial component of aging and its study will likely yield key insights into the aging process [2]. In this line, the connection between epigenetics and aging has recently been reinforced by the construction of epigenetic clocks based on the analysis of age-associated changes in DNA methylation [3-7]. Importantly, while these clocks predict age with high accuracy, their link to cellular function has not been established yet [8]. This is likely to occur due to the fact that DNA methylation is only one of several mechanisms that affect chromatin states and transcription. On the other hand, transcriptomics and proteomics clocks although less accurate have succeeded at providing a clear link to cellular function [9-16]. However, their readout does not allow for a simple understanding of how underlying epigenetic mechanisms change with age. For this reason, we aimed to develop a novel aging clock that provides insight into epigenetic mechanisms while also being biologically relevant.

Changes in chromatin accessibility, resulting from the cumulative effect of multiple epigenetic mechanisms, have been shown to occur during aging [17-21]. Multiple studies have identified associations between aging and loss of repressive epigenetic marks, thus proposing the “loss of heterochromatin” theory of aging [22]. We envision that these changes could allow us to predict age and at the same time increase our understanding of epigenetic dysregulation. Towards this goal, we generated chromatin accessibility and transcriptomic profiles from human blood samples spanning a broad range of ages using ATAC-seq [23] and RNA-seq respectively. We then analyzed age-related changes in accessibility and their effect of transcription. Subsequently, we used an elastic net regression model to predict age from chromatin accessibility profiles with good accuracy. Finally, we characterized this novel aging clock by investigating its features and comparing its performance to that of transcriptome-based clocks.

## Results

### Profiling human blood samples over a wide age range

Blood samples were acquired from 159 healthy donors (117 men, 42 women) covering an age range from 20 to 74 years (Figure 1). Peripheral blood mononuclear cells (PBMCs) were isolated to generate ATAC-seq profiles from 157 samples, of which 141 (104 men, 37 women) passed quality controls. A representative histogram of fragment size distribution is included in Supplementary Figure 1c. From these samples, we detected a total of 42557 open chromatin regions (OCRs), of which 34.1% lay within 1 kbp of transcription start sites (TSS) and were thus annotated as promoters, 50.7% contained sites with reported enhancer activity, 5.6% were annotated as both promoters and enhancers. The remaining 9.6% OCRs which did not lie in the proximity of TSSs and had no reported enhancer activity will be referred to as “unannotated” (Figure 2b). Principal component analysis of accessibility profiles partially separated samples by age (Supplementary Figure 1a). Additionally, we performed RNA-seq on all samples, from which we detected the expression of 21828 genes. Of these samples, 151 passed quality control and 135 had a matching ATAC-seq sample. Principal component analysis of expression profiles partially separated samples by age (Supplementary Figure 1a). Finally, we used flow cytometry to measure the proportions of monocytes, granulocytes, lymphocytes, total T cells, CD4+ T cells, CD8+ T cells, B cells, and NK cells in all samples (Supplementary Figure 1b). During aging we detected an increase in the proportions of NK cells (Pearson’s r = 0.31, p < 0.001) and a decrease in the numbers of total T cells (Pearson’s r = -0.22, p = 0.005) and CD8+ T cells (Pearson’s r = - 0.24, p = 0.002). The proportions of monocytes, granulocytes, lymphocytes, CD4+ T cells, and B cells did not significantly correlate with age. Similar changes in PBMC compositions have been reported in previous studies [19,24,25].

**Figure 1.**
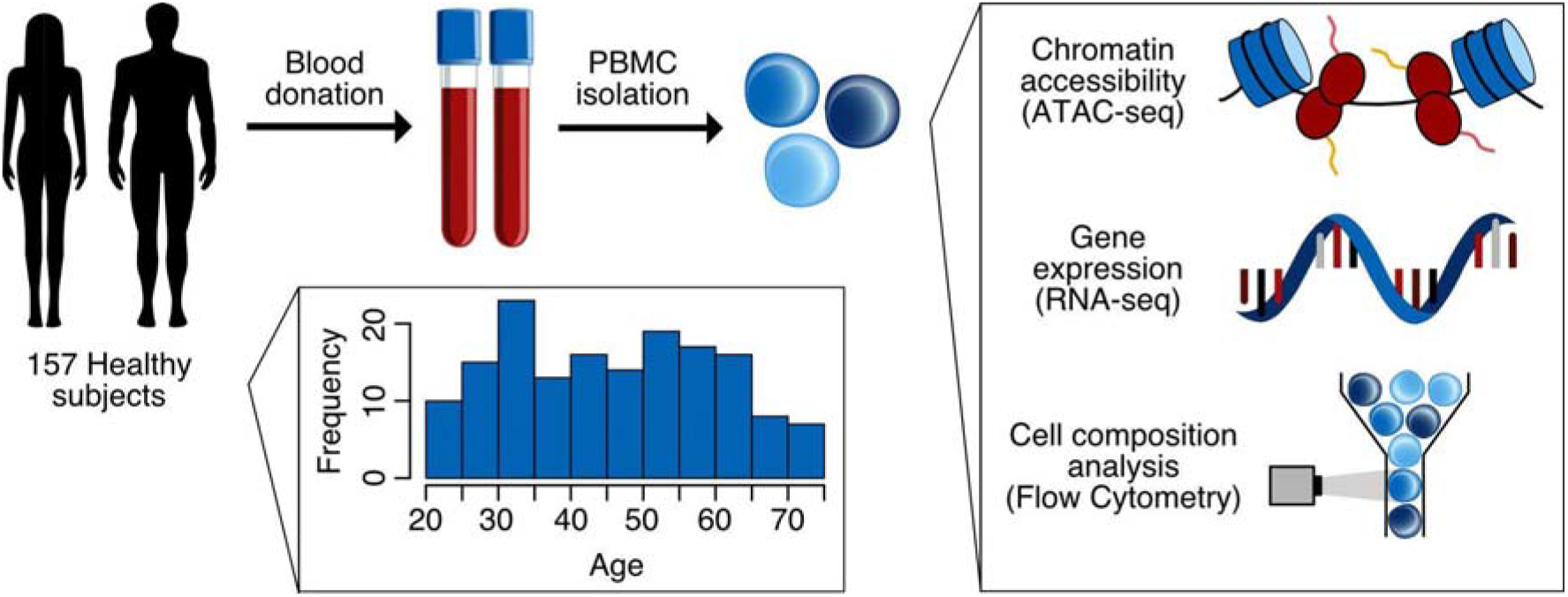
Schematic outline of sample acquisition. PBMCs were isolated from blood samples of 157 healthy donors (ages 20 – 74) with a nearly uniform distribution. ATAC-seq and flow cytometry profiles were generated from all samples, whereas RNA-seq data was generated on a subset of samples.

**Figure 2.**
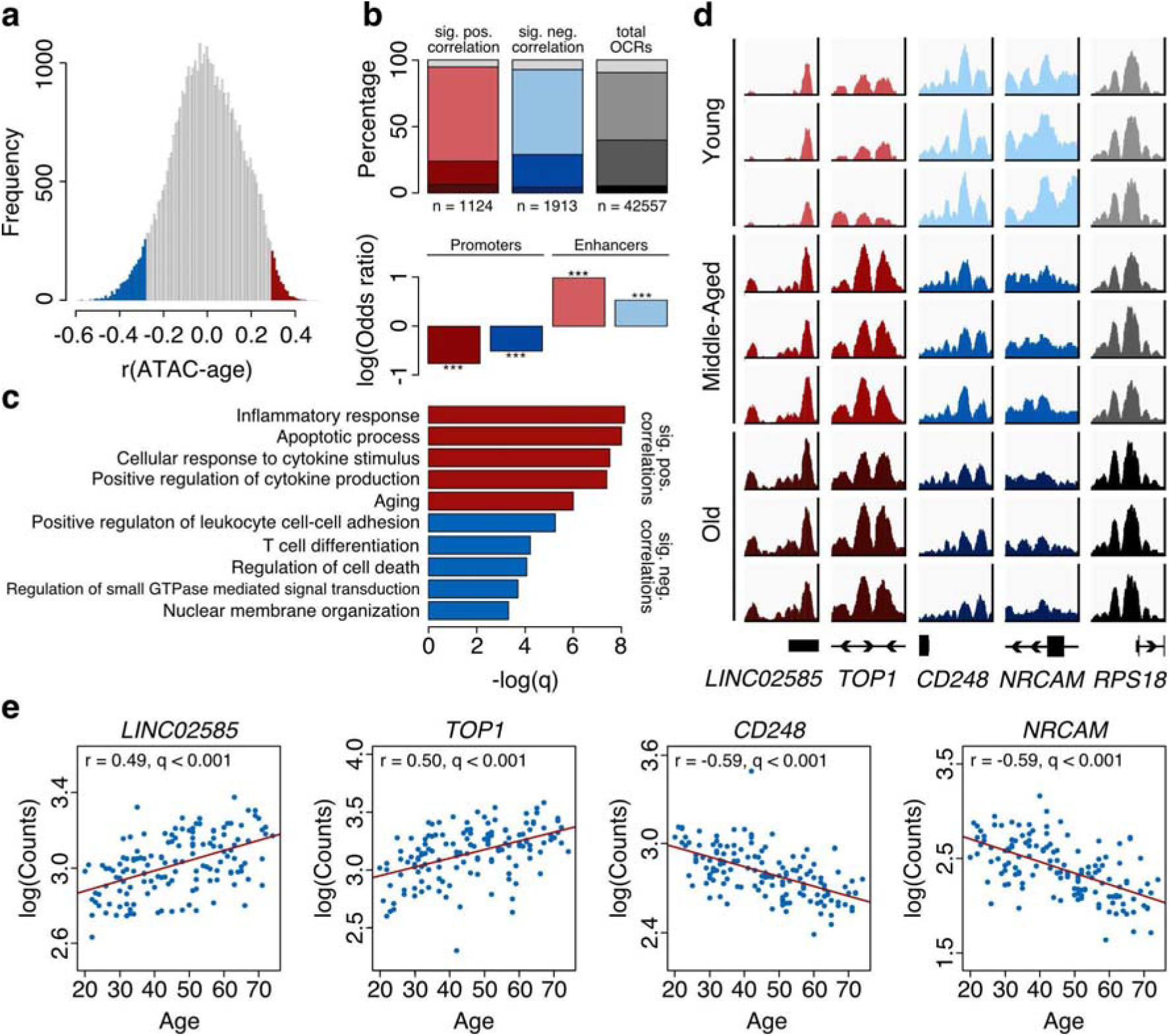
Chromatin accessibility changes with age. **a** Distribution of correlations between chromatin accessibility and age (spearman r). Statistically significant closing features are highlighted in blue, while statistically significant opening features are highlighted in red (FDR < 0.01). **b** Annotation of statistically significant features to regulatory elements. Enrichment of promoters and enhancers compared to all features (background). Log(odds ratios) and p-values were calculated using Fisher’s Exact test (*** = p < 0.001). **c** Gene Ontology (GO) of significantly correlated OCRs. GO terms are plotted against the -log of FDR corrected p-values (q-value). Opening OCRs are indicated in red, whereas closing OCRs are indicated in blue. The top five GO terms from the highest hierarchy (most specific) are shown. **d** The top two age-correlating opening and closing OCRs are shown. The last column represents a housekeeping gene that did not correlate with age. Each row represents one sample, whereas the young group includes samples from donors aged 20 - 32 years, the middle-aged group includes samples from donors aged 41 - 53 years and the old group includes samples from donors aged 62 - 74 years. **e** Accessibility vs age scatterplots of the top four OCRs.

### Chromatin accessibility changes in a site-specific manner during aging

To better understand the effect of aging on the epigenome, we analyzed global and site-specific changes in chromatin accessibility. Although based on the heterochromatin loss theory of aging we might expect to see a gradual de-repression of chromatin outside our OCRs, we did not detect any significant correlation between the fraction of reads within peaks (FRIP) and age (Pearson’s r = -0.13, p = 0.14, Supplementary Figure 2a). Next, we investigated whether we could observe a change in accessibility profiles during aging, such as an increase in peak width as has been reported for ChIP-seq data of specific histone marks [26]. Thus, we compared the average coverage profile around transcription start sites (TSS) of samples from young (< 35 years old, n = 40) and old (> 55 years old, n = 44) donors but found no significant change (Supplementary Figure 2b). As for site-specific changes, we observed a consistent opening of chromatin with age in 1124 OCRs and closing in 1913 OCRs (Spearman’s r, FDR < 0.01) (Figure 2a). Several examples of coverage profiles for OCRs that open, close, or do not change with age are shown in Figure 2d, whereas Figure 2e shows the correlation between the accessibility of the same OCRs and age. Among the opening OCRs, 6.3% were annotated as both promoters and enhancers, 17.5% as promoters, 70.9% as enhancers, and 5.3% were unannotated. Among the closing OCRs, 4.4% were annotated as promoters and enhancers, 24.6% as promoters, 63.7% as enhancers, and 7.3% were unannotated. Interestingly, we observed a statistically significant enrichment of enhancers and a statistically significant depletion of promoters in both the opening and the closing OCRs compared to the background (Fisher’s Exact Test, Figure 2b). This suggests that enhancers could be particularly sensitive to changes in accessibility during aging. Next, we linked OCRs to genes and investigated their involvement in biological processes. For this purpose, we associated promoters to their closest gene, whereas we assigned enhancers to genes using the PEREGRINE dataset: a collection of enhancer-gene links predicted based on ChIA-PET, eQTL, and Hi-C of multiple tissues, including blood [27]. Interestingly, the opening OCRs were linked to 1541 genes, 1388 of which were expressed. On the other hand, the closing OCRs were linked to 1891 genes, 1693 of which were expressed. We used the genes that were expressed to perform gene ontology (GO) analysis (Figure 2c). The gene set associated with the opening OCRs revealed an enrichment of genes associated with inflammation, apoptosis, and aging. Similarly, the gene set associated with the closing OCRs yielded associations with cell-cell adhesion, cell differentiation, cell death, GTPase-mediated signal transduction, and nuclear membrane organization. In conclusion, we found that chromatin accessibility of PBMCs does not undergo global changes during aging, at least in the age range that we analyzed (20-74). Instead, we detect changes in specific regulatory elements, most commonly enhancers, which are associated with inflammation and innate immunity. The correlations between accessibility at specific OCRs and age suggest that it should indeed be possible to construct an aging clock based on chromatin accessibility similarly to clocks based on DNA methylation.

### Changes in chromatin accessibility relate to changes in expression

One limitation of aging clocks based on DNA methylation is that changes in methylation do not correlate with robust changes in the expression of downstream genes [28]. Therefore, we asked whether in our dataset we were able to observe changes in expression coherent with changes in chromatin accessibility. GO analysis revealed that genes upregulated during aging were associated with inflammatory pathways such as tumor necrosis factor (TNF) and nuclear factor-kB (NF-kB), while several of the terms related to downregulated genes suggested a decline in specific immune functions related to B cell function and phagocytosis (Figure 3a). Investigating in more detail, we identified several genes whose expression and accessibility at regulatory elements both correlated to age (Figure 3b, Figure 3c shows coverage plots of one such gene: CD248) and sought to determine if these were more common than expected by chance. Therefore, we compared the age correlations of genes linked to OCRs with positive correlation with age (Spearman r > 0, FDR < 0.01), negative correlation with age (Spearman r < 0, FDR < 0.01) and no correlation with age (FDR > 0.01) (Figure 3d). We found that overall, genes linked to promoters whose accessibility increased with age were upregulated during aging (Kolmogorov-Smirnov Test, D = 0.33, p < 0.001), similarly genes linked to promoters which closed with age tended to be downregulated in aging (Kolmogorov-Smirnov Test, D = 0.38, p < 0.001, Figure 3d). Interestingly, this pattern was much weaker when we looked at enhancers whose accessibility increased with age (Kolmogorov-Smirnov Test, D = 0.07, p < 0.001) and enhancers whose accessibility decreased with age (Kolmogorov-Smirnov Test, D = 0.09, p < 0.001). In other words, genes linked to these enhancers only showed a small bias towards the direction of change of their enhancer (Figure 3d). This difference could have several explanations. First, it should first be mentioned that the enhancer-gene links in the PEREGRINE dataset were described in a variety of tissues and might not be optimal for the analysis of PBMCs where some interactions might be missing. To verify this, we repeated the analysis restricting to enhancer-gene links that have been verified in whole blood, however, the outcome did not change (Supplementary Figure 2c). It is also possible that accessibility changes at enhancers could be less important for transcriptional control, either because they could be overshadowed by changes at the promoter or because transcriptional regulation at enhancers could occur by mechanisms that do not alter chromatin accessibility directly. In this line, it has been shown that regions marked with the repressive histone mark H3K27me3 can remain accessible to binding by general transcription factors [29-31].

**Figure 3.**
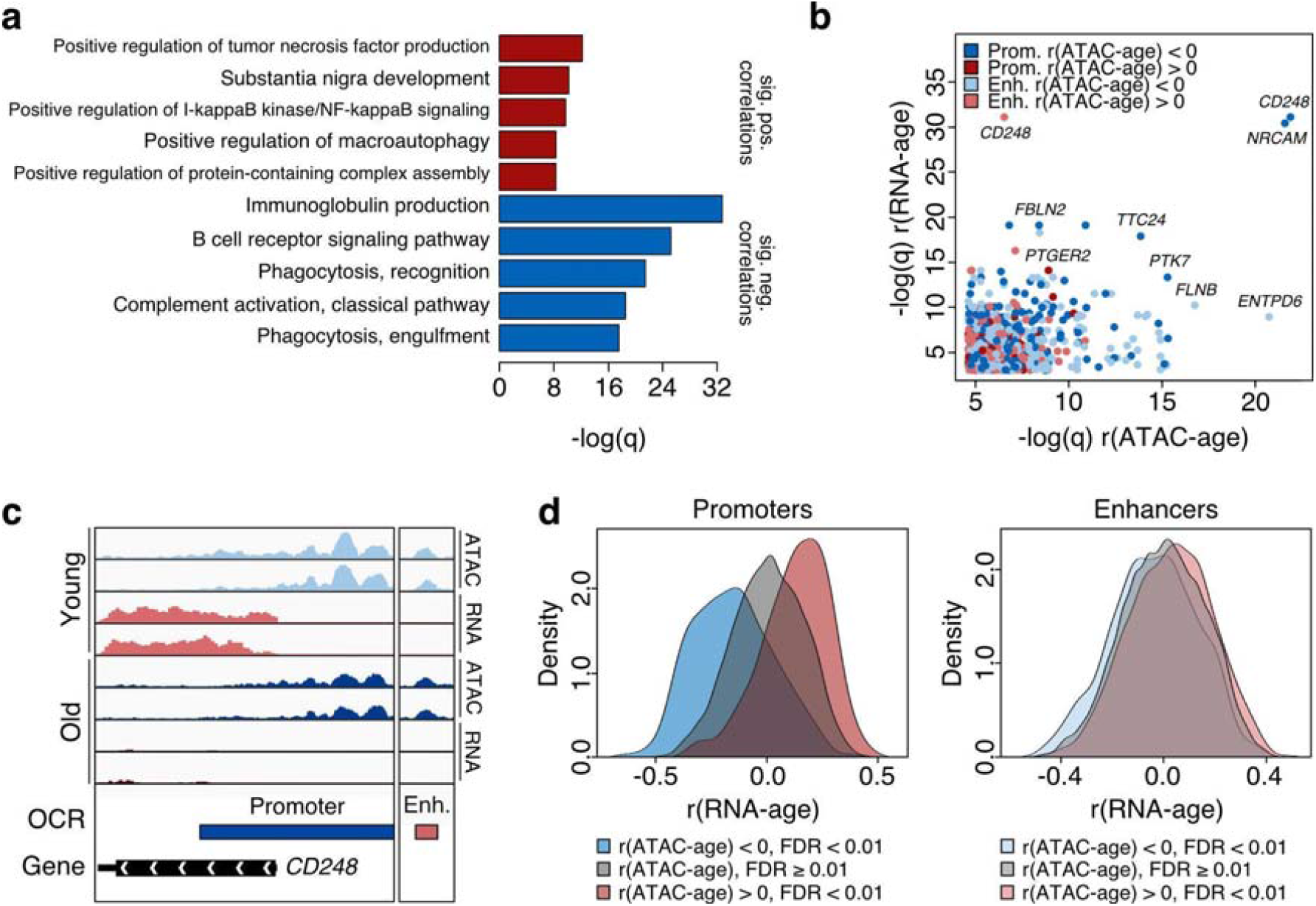
The relationship between accessibility and expression. **a** Gene Ontology (GO) of genes whose expression significantly correlates with age. GO terms are plotted against the -log of FDR corrected p-values (q-value). More expressed genes are indicated in red, whereas less expressed genes are indicated in blue. The top five GO terms from the highest hierarchy (most specific) are shown. **b** Genes whose expression and accessibility at regulatory elements correlated with age. The x-axis represents the significance of correlation in the ATAC-seq data while the y-axis represents the significance of correlation in the RNA-seq data. **c** ATAC-seq and RNA-seq coverage tracks for the gene CD248, whose expression and accessibility at promoter and enhancer both decrease with aging. Two young and two old samples are shown. **d** Distribution of correlation between transcription levels of genes and age linked to promoters/enhancers whose accessibility is positively correlated with age (r > 0, FDR < 0.01), negatively correlated with age (r < 0, FDR < 0.01) or not correlated with age (FDR ≥ 0.01).

### Chromatin accessibility predicts age

Having found many site-specific changes in chromatin accessibility with age, we next investigated whether these changes could be used to predict the age of the blood donors. To do so, we trained an elastic net regression model on the 141 ATAC-seq samples which passed quality control. We used nested cross-validation to tune hyperparameters and estimate the performance of the model. Across the outer folds of the nested cross-validation, the model selected 183 ± 58 OCRs as predictors and predicted age with an RMSE of 7.69 ± 1.67, MAE of 5.69 ± 1.59 and r of 0.83 ± 0.08 (Figure 4a). A final model trained with the whole dataset selected a total of 297 OCRs, 139 of which were taken with a positive coefficient and 158 with a negative one. Of all the OCRs selected by the model, 3.7% were annotated as promoters and enhancers, 29.6% as promoters, 55.2% as enhancers, and 11.4% were unannotated. Interestingly, clock sites were slightly depleted of promoters compared to background (Fisher’s Exact Test: log(odds ratio) = -0.28, p = 0.02), but they were not enriched of enhancers (Fisher’s Exact Test: log(odds ratio) = 0.10, p = 0.38). This contrasts with the finding that age-correlated OCRs are enriched in enhancers. As elastic net models do not simply select the features that best correlate to the response variable but aim to eliminate redundant features, a possible explanation is that many enhancers could be strongly correlated with each other and thus are not chosen by the model. We then analyzed the relationship between the accessibility of OCRs selected by the clock and gene expression. As expected, we found that the age correlation of promoters correlated with the age correlation of the linked genes (Pearson’s r = 0.47, p < 0.001, Figure 4b), whereas this relationship was weaker in enhancers but still significant (Pearson’s r = 0.15, p = 0.02, Figure 4b). This suggests that a biologically meaningful subset of OCRs selected as predictors by the clock might drive changes in transcription that are coherent with their changes in accessibility during aging, strengthening the causal link between the clock and physiological changes. With this knowledge, we investigated the potential biological role of the OCRs selected by the clock. GO analysis of clock sites did not yield any terms, presumably because elastic net regression removes redundant predictors, meaning it is unlikely that multiple predictors will be chosen within the same biological process. The same held true when we performed GO analysis separately for promoters and enhancers. Instead, we manually investigated the clock OCRs with the 20 largest absolute coefficients in the model (Figure 4c). One of these OCRs was the promoter of *GREM2*, a gene that encodes a senescence-associated secretory phenotype (SASP) factor with known association with aging in adipose tissue and skin [32,33]. This OCR was also selected in every nested cross-validation model, highlighting its robustness to predict age. In our data, chromatin accessibility at the *GREM2* promoter opens with age (r = 0.44, q < 0.001) and we observed an increase of *GREM2* transcripts with age (r = 0.40, q < 0.001). *IKZF2*, also among the top 20 genes, encodes for the transcription factor Helios, which is mainly expressed in hematopoietic stem and progenitor cells and represses transition towards the myeloid lineage. *IKZF2* null mice unsurprisingly have a shift from the lymphoid towards the myeloid lineage, which is also observed in aged animals [34]. In our data, chromatin accessibility at the *IKZF2* promoter strongly decreases with age (r = -0.51, q < 0.001), however *IKZF2* transcripts do not significantly correlate with age (r = -0.08, q = 0.60).

**Figure 4.**
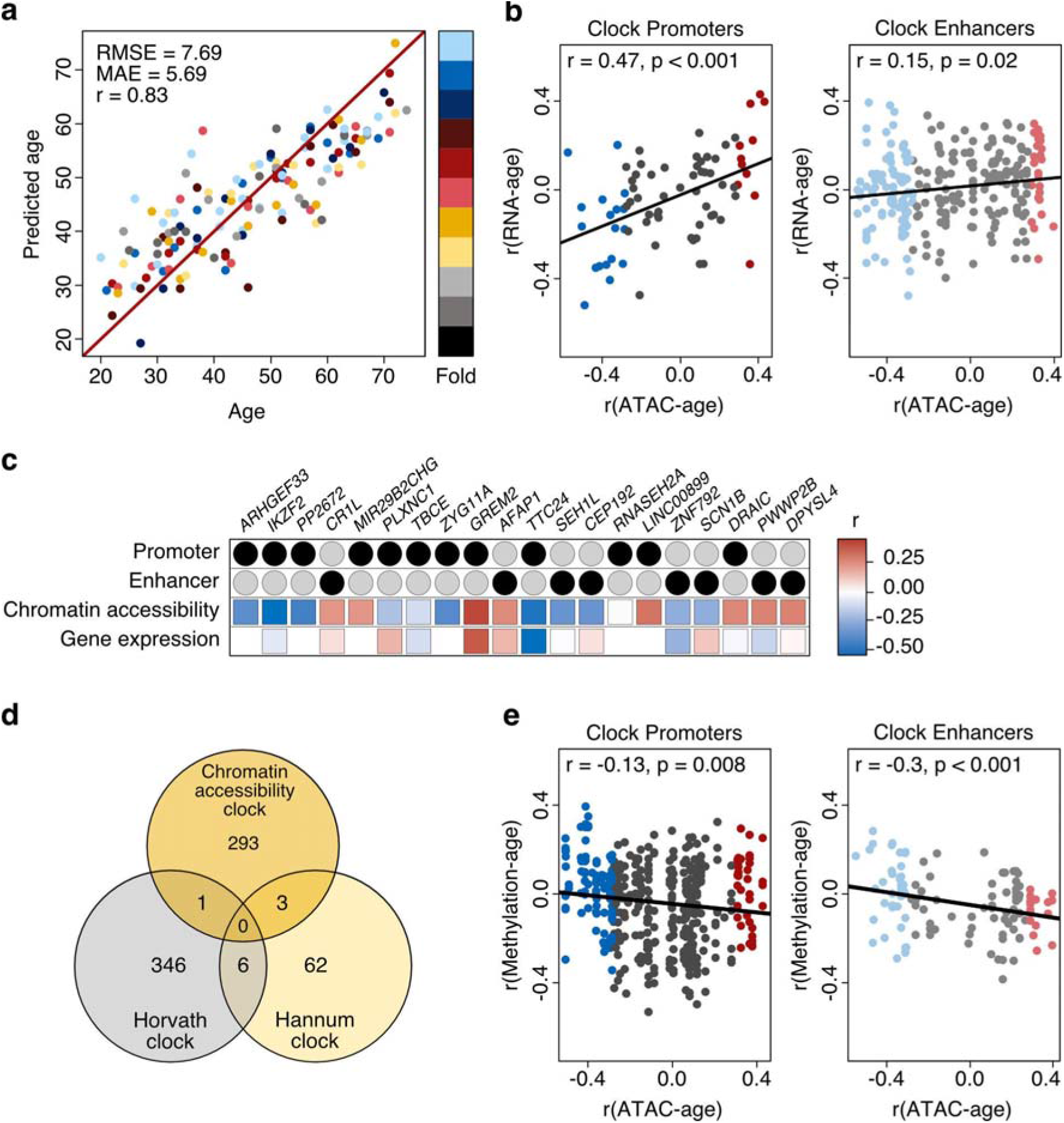
Chromatin accessibility predicts age. **a** Development of an age predictor using nested-cross validation with 11 outer folds (11 different models, each sample in the test set once). The scatter plot shows the test set data from each fold. The x-axis represents the real age, and the y-axis represents the predicted age. The average root mean squared error (RMSE), the average median absolute error (MAE), and the average Pearson correlation coefficient (r) are shown. **b** Age correlation of gene transcription vs age correlation of linked clock promoters/enhancers. **c** Annotation of the top 20 OCRs of the final model ranked by their absolute coefficient. Correlation in chromatin accessibility and correlation in expression level with age are indicated for each OCR/gene pair. **d** Intersection between genomic sites used by the chromatin accessibility clock, the Horvath clock, and the Hannum clock. **e** Age correlation of methylation sites from the Hannum dataset vs age correlation of linked clock promoters/enhancers.

### The chromatin accessibility clock relies on a different aging signature compared to methylation clocks

Next, we looked for similarities between our clock and previously published methylation-based aging clocks (Figure 4d). The intersection of the chromatin accessibility clock OCRs with the 353 CpG sites selected by the Horvath clock [5] identified an overlap in the promoter of *UBE2V1*. This site had a positive coefficient in the Horvath clock and a negative one in our chromatin accessibility clock, which is coherent with the role of DNA methylation in establishing heterochromatin. At the RNA level, however, we did not find a correlation between *UBE2V1* transcription and age. The intersection of the chromatin accessibility clock OCRs with the 71 CpG sites selected by the Hannum clock [6] identified three overlapping features. One is in the promoter of *MIR29B2CHG* and enhancer of *CR1L*, one is in the promoter of *ARHGEF33*, and one is in the enhancer of *KLF13*. Excitingly, we note that OCRs related to MIR29B2CHG, CR1L, and ARHGEF33 were among the 20 most important predictors for our clock. The coefficients related to *ARHGEF33* and *KLF13* were taken by our clock and the Hannum clock with opposite signs, however, for the site related to *MIR29B2CHG* and *CR1L* this was not the case. At the RNA level, transcription of *ARHGEF33* and *MIR29B2CHG* was not detected in our dataset. *CR1L* and *KLF13* on the other hand are transcribed, but their transcription levels do not correlate with age. Although few of our accessibility clock’s sites contained CpG sites used by the Hannum and Horvath clocks, we investigated whether the age-related changes in accessibility observed in our data could be attributed to changes in DNA methylation. For this, we focused on the Hannum dataset as it was generated from blood samples. Of the 485577 CpG markers in the Illumina Infinium 450 Human Methylation array, 141536 fell within our OCRs (107097 in promoters, 13191 in enhancers, 14782 in OCRs annotated as both promoters and enhancers, and 2679 in unannotated OCRs). We found that enhancer opening during aging broadly correlated with methylation decreasing and vice versa (Pearson r = -0.30, p < 0.001, Figure 4e). However, this pattern was much weaker in promoters (Pearson r = -0.13, p < 0.008). These results indicate some degree of connection between accessibility and methylation data in blood. However, given that the majority of CpG sites in the Infinium 450 array are located within promoters where this relationship is weak, we believe that our chromatin accessibility clock and published methylation clocks predict age based on distinct epigenetic signatures. However, matched methylation and ATAC-seq samples would be required to draw final conclusions.

### Changes in cellular composition alone do not explain the clock’s performance

To further characterize the clock and because our flow cytometry data revealed a correlation between the size of certain immune cell populations and age, we wanted to understand to what extent the performance of the chromatin accessibility clock depends on changes in cell composition as opposed to cell-intrinsic changes in accessibility. Importantly, a clock trained on cell composition alone performed significantly worse than the chromatin accessibility clock: RMSE 13.56 ± 1.64, MAE 10.34 ± 1.66, r 0.39 ± 0.15 (t-Test: p-values < 0.001, Supplementary Figure 3a, b). Similarly, when we trained the chromatin accessibility clock providing cell composition as potential covariates, none of the 11 models selected by nested cross-validation used cell composition features as predictors. These findings suggest that age-related changes in cell composition are not sufficient to predict age accurately. Nonetheless, it is possible that measuring further subpopulations such as naive, activated, and exhausted T cells could improve the performance of a clock based on cell composition.

### Gender has little influence on the clock’s performance

Previous publications have reported important differences in immune aging between men and women [19]. We note that in our chromatin accessibility clock, the error on predictions did not differ significantly between men and women (t-Test: p = 0.29, Supplementary Figure 3c). Nonetheless, we sought to determine whether accounting for gender differences could improve clock performance. As for the case of cellular composition, the gender covariate was never picked by the models we trained. Comparison of accessibility clocks trained on only male samples (n = 74) and clocks trained on mixed-gender samples (m = 37, f = 37) revealed that clocks trained on only male samples (RMSE 9.76 ± 3.05, MAE 7.08 ± 2.86 and r 0.75 ± 0.12) did not perform significantly better than those trained on mixed samples (RMSE 9.45 ± 1.61, MAE 7.28 ± 2.1 and r 0.71 ± 0.18) (t-Test: p-values = 0.47 (RMSE), 0.85 (MAE), 0.58 (r), Supplementary Figure 3d, e). While there are undeniably differences in epigenetic aging trajectories between genders, a fraction of these changes are conserved [19]. Given that the clock only selects a small subset of all age-related OCRs, there are likely to be plenty of non-gender-specific predictive sites that it can choose from, thus making correction for gender seemingly unnecessary.

### Changes in chromatin accessibility predict age better than changes in gene expression

Next, we wanted to compare the predictive power of our aging clock based on chromatin accessibility with that of clocks based on gene expression, therefore, we used samples from donors in which we obtained both ATAC-seq profiles and RNA-seq profiles to construct two separate clocks (Figure 5a). In this direct comparison, the predictions of the two clocks correlated strongly with each other (Pearson’s r = 0.80, p < 0.001) however the chromatin accessibility clock performed significantly better by two metrics (RMSE 7.36 ± 1.38, MAE 5.66 ± 1.35 and r 0.85 ± 0.06 for the chromatin accessibility clock compared with RMSE 9.48 ± 1.94, MAE 7.22 ± 2.06 and r 0.73 ± 0.12 for the gene expression clock, t-Test: p-values = 0.008 (RMSE), 0.051 (MAE), 0.008 (r), Figure 5b). Thus, in our dataset, accessibility features appear to allow for better age prediction than gene expression features. However, we note that the BiT age clock was able to obtain better performance from gene expression data by binarizing the features in a dataset with a comparable number of samples: n = 131, RMSE 8.41, MAE 5.24, r 0.96 [16]. The large difference in r score despite similar RMSE and MAE is likely to depend on the standard deviation of age in the dataset: specifically, it was shown that the standard deviation of age in a dataset positively correlates with the r score [5]. In conclusion, it appears that changes in chromatin accessibility may offer a better ability to predict age when compared to changes in transcription. Nonetheless, more samples, possibly from different tissues, would be required to confirm this result.

**Figure 5.**
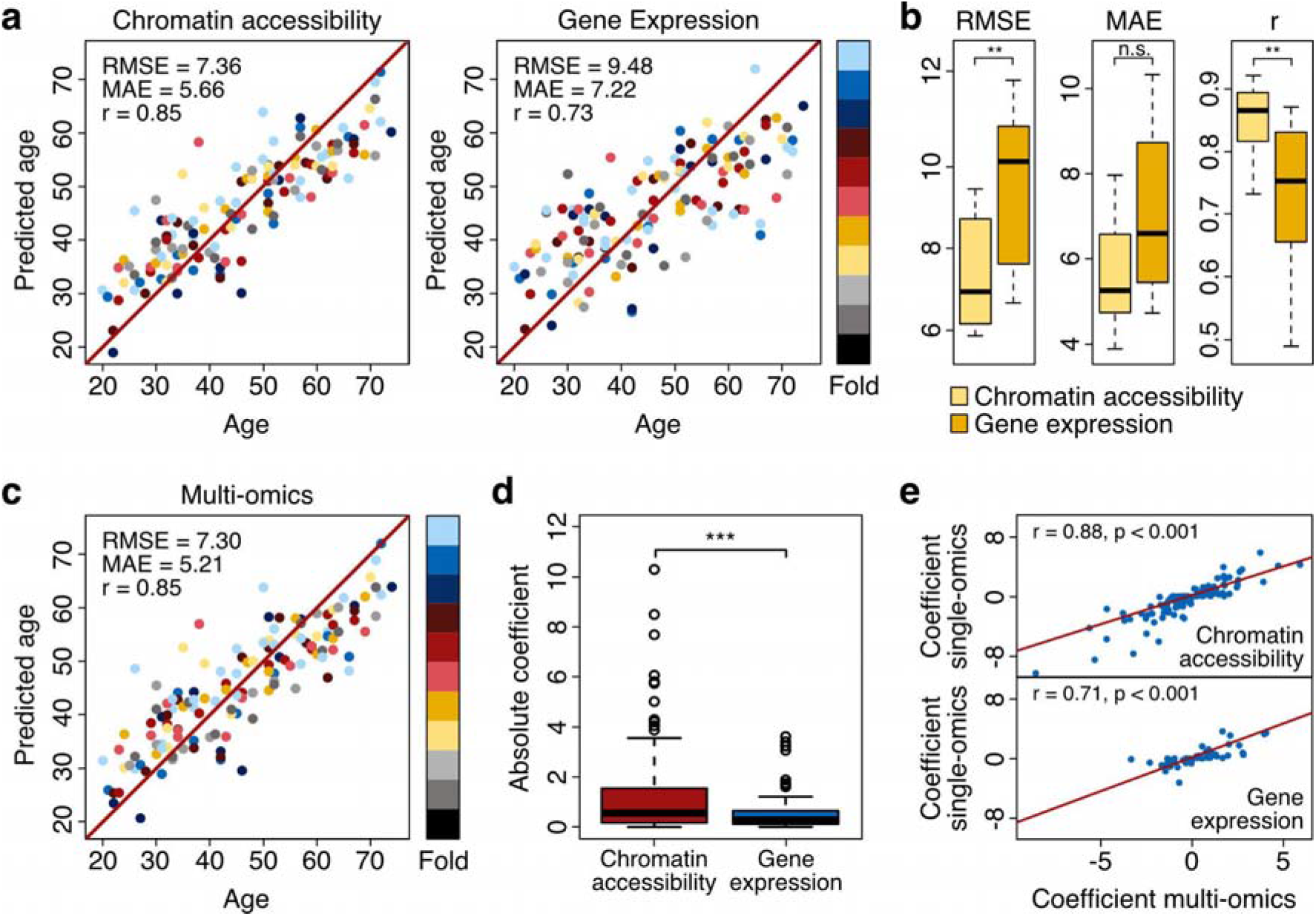
Chromatin accessibility predicts age better than gene expression. **a** Comparison of chromatin accessibility and transcriptomics clocks trained on matched ATAC-seq and RNA-seq samples (n = 135). **b** Comparison of scores of the chromatin accessibility and transcriptomics clocks. **c** Multi-omics clock trained on ATAC-seq and RNA-seq features. **d** Comparison of selected ATAC-seq and RNA-seq features in terms of absolute coefficients. **e** Correlation between coefficients in the multi-omics and single-omics clock.

### Multi-omics clock based on chromatin accessibility and gene expression changes

Finally, we concatenated the ATAC-seq and RNA-seq profiles from the matching samples (n = 135), using a total of 64385 features (42557 ATAC-seq and 21828 RNA-seq) as input to the elastic net model. This multi-omics clock predicted age with an RMSE of 7.3 ± 1.4, MAE of 5.21 ± 1.64, and r of 0.85 ± 0.06 (Figure 5c), a similar performance to the results achieved by the chromatin accessibility clock estimated on the same set of samples (Figure 5a). A final model trained with the whole multi-omics dataset selected a total of 232 features, from which 158 corresponded to ATAC-seq features and 74 corresponded to RNA-seq features, preserving the initial ATAC to RNA features proportion. To examine the influence of the chromatin accessibility and gene expression features on the final model, we compared the absolute values of the coefficients assigned to the selected ATAC-seq and RNA-seq features (Figure 5d). We found that the chromatin accessibility features exert a significantly stronger influence than the gene expression features on the multi-omics clock, detecting a significant difference between the absolute coefficients of the ATAC-seq and RNA-seq covariates (t-Test, p < 0.001). We also analyzed the overlap between the features selected by the multi-omics clock and each of the two single-omics models fitted on the set of matching samples. On the one hand, among the 351 OCRs finally selected by the chromatin accessibility clock, a significant overlap of 128 ATAC-Seq features selected by the multi-omics clock was found (Fisher’s Exact test, p < 0.001). On the other hand, from the set of 351 RNA-seq features selected by the gene expression clock, we detected a significant overlap of 52 RNA-Seq features selected by the multi-omics clock (Fisher’s exact test, p < 0.001). Additionally, we examined the relationship between the coefficients of the overlapping features assigned by the model in the multi-omics and single-omics settings. For both the chromatin accessibility and gene expression features, we saw a significant correlation between the coefficients in the multi-omics and single-omics models (Pearson’s r = 0.88, p < 0.001 for the ATAC-Seq features, and Pearson’s r = 0.71, p < 0.001 for the RNA-Seq features) (Figure 5e), showing that the overlapping features influence both the single-omics and multi-omics clocks in an similar manner. In conclusion, a combination of chromatin accessibility and gene expression features can also predict age accurately, suggesting the existence of a multi-omics aging signature.

## Conclusions

In this study, we have generated ATAC-seq, RNA-seq, and flow cytometry data from PBMCs isolated from healthy individuals in the age range of 20 - 74 years. We analyzed changes in chromatin accessibility during aging and detected the opening of specific sites related to inflammation, as well as the closing of sites linked to adaptive immunity. We then investigated the link between gene transcription and chromatin accessibility at related regulatory elements. In general, age-related changes in transcription and accessibility were related to similar biological processes, and at a site-specific level, we found that accessibility changes at promoters drove coherent transcriptional responses. However, enhancer accessibility seemed to have a much weaker effect on the transcription of related genes, suggesting a possible difference in regulatory mechanisms between promoters and enhancers which could not be captured by ATAC-seq. We then showed for the first time that chromatin accessibility profiles of PBMCs can be used to predict the age of donors, with an RMSE of 7.69 years, MAE of 5.69 years, and r of 0.83. This clock predicts age accurately, although we expect performance could be improved even further by addition of more samples to the training set. Next, we compared the performance of clocks trained on matching ATAC-seq and RNA-seq profiles and found that our chromatin accessibility clock performed significantly better than its transcriptomic counterpart. Finally, we developed the first multi-omics clock based on chromatin accessibility and transcriptome features. This multi-omics clock performed similarly to the accessibility clock and relied on accessibility features more than on gene expression features, once again suggesting that chromatin accessibility data may allow for better age prediction than transcriptomic data. In conclusion, we demonstrated the possibility of constructing an aging clock based on chromatin accessibility that retains a direct link to age-associated alterations in epigenetic states, while at the same time provides insights regarding changes in cellular function that occur with age.

## Acknowledgments

The study was supported by the Novartis Foundation for Medical-Biology Research.

## Authors Contributions

C.R. and F.M. wrote the manuscript draft. C.R. isolated PBMCs, prepared ATAC-seq libraries, extracted RNA, analyzed cell composition, and carried out bioinformatics analyses. F.M. developed the chromatin accessibility clock, carried out bioinformatics analyses, and contributed to blood sample processing. K.P. pre-processed the RNA-seq data. V.P. contributed to pre-processing of the ATAC-seq data. Both K.P. and V.P. contributed to bioinformatics analyses. G.L.G. developed the multi-omics clock. L.H. contributed to the ATAC-seq setup. F.v.M. provided reagents and advised on the study and analysis. A.O. designed and supervised the study. All authors contributed to reviewing and editing the manuscript.

## Methods

### Blood collection

Anonymized whole blood from 157 donors between the ages of 20 and 74 was obtained from the Interregional Blood Transfusion center in Lausanne-Epalinges, Switzerland. The internal review board approved the study, and all donors gave written consent to the usage of their blood for research purposes. Samples were processed within 4.5 hours after blood collection.

### PBMC Isolation

Blood was diluted with equal amounts of Dulbecco’s phosphate-buffered saline (Gibco) and layered on top of Histopaque-1077 (Sigma-Aldrich). Density gradient centrifugation was carried out according to the manufacturer’s protocol and PBMCs were collected and washed. Cells were counted on a LUNA-II Automated Cell Counter (Logos Biosystems) and immediately aliquoted for ATAC-Seq library preparation, RNA extraction, and PBMC staining/fixation. All protocols were carried out simultaneously.

### ATAC-Seq library preparation

ATAC-Seq library preparation was performed according to the Omni-ATAC protocol [35] using Tn5 provided by the EPFL Protein Production and Structure facility. Transposed fragments were purified using the MinElute PCR Purification Kit (Qiagen). The eluate was PCR amplified using 2x NEBNext Master Mix (NEB) and pre-mixed primers with unique dual indexes for Illumina sequencing (IDT). The library was purified by double-sided bead size selection using SPRIselect (Beckman Coulter).

### RNA extraction

RNA extraction was performed using the Monarch Total RNA Miniprep Kit (NEB) according to the manufacturer’s protocol.

### PBMC staining and flow cytometry

Cells were stained with Ghost-Dye/V510, CD3+/APC, CD4+/FITC, CD8+/APC-Cy7, CD16+/PE, CD19+/PE-Cy7 and CD56+/APC (Biolegend). Cells were fixed in Fixation and Permeabilization Solution (BD). A Cytoflex S flow cytometer (Beckman Coulter) was used to analyze the subpopulation ratios.

### ATAC sequencing and data pre-processing

ATAC-Seq libraries were subjected to 150 bp paired-end sequencing on an Illumina NovaSeq 6000 by Novogene (UK) Company Limited with a sequencing depth of 30 million reads. Raw reads were adapter and quality trimmed using Trim Galore! [36] and mapped to the GRCh38 build of the human genome using bowtie2 (with settings --very-sensitive -k 10 - X 1000 --dovetail) [37]. Before peak calling, raw bams were filtered to remove reads with multiple mappings, PCR duplicates, and mitochondrial reads using samtools [38] and Picard tools [39]. Peaks were called using Genrich [40] in ATAC mode (-j) with a maximum q value of 0.05 (-q 0.05) and ignoring reads overlapping the ENCODE blacklist [41]. To define a common peakset, the union of the peak sets of individual samples was computed using BEDTools merge [42]. Subsequently, counts representing the accessibility at each peak were computed for every sample by intersecting regions centered around Tn5 cut sites as output by Genrich with the common peakset using bedtools intersect. Counts were first transformed to read densities by dividing the counts at each peak by the length of the peak in kilobases and then normalized by dividing by the total number of reads-in-peaks in millions.

Low-quality samples and outliers were detected using the elliptic envelope method on the first two principal components of the normalized counts in linear scale [43]. This removed 16 samples, including all samples with insufficient sequencing depth.

Coverage bigWig tracks were generated from regions centered around Tn5 cut sites using deepTools bamCoverage [44] with the same scaling factors used to normalize the counts. TSS profiles were generated from the bigwig tracks using deeptools computeMatrix and plotProfile.

### RNA sequencing and data pre-processing

RNA-Seq library preparation and sequencing was performed by Novogene (UK) Company Limited on an Illumina NovaSeq 6000 in 150 bp paired-end mode. Raw FASTQ files were assessed for quality, adapter content and duplication rates with FastQC. Reads were aligned to the Human genome (GRCh38) using the STAR aligner (v2.7.9a) [45] with ‘-- sjdbOverhang 100’. Number of reads per gene was quantified using the featureCounts function in the subread package [46]. Ensembl transcripts were mapped to gene symbols using the mapIds function in the AnnotationDbi package [47] with the org.Hs.eg.db package [48]. Raw counts were normalized using the trimmed means of M-values (TMM) method with EdgeR [49] and filtered to remove genes that did not reach 10 counts in at least 10 samples. Finally, 5 outliers were removed using the same strategy employed for the ATAC-seq dataset.

### Clock Construction and Characterization

Training and validation of the elastic net model were carried out in Python using the Scikit-learn module [43]. Samples were assigned to 11 groups so that the age composition in each group would cover the age range uniformly. Nested cross-validation was used to tune hyperparameters and estimate the performance of the model. Both the outer and inner cross-validation loops were run as leave-one-group-out cross-validation, meaning that the outer loop used each of the 11 groups once as a test set, while the inner loop alternated over the remaining 10 groups. Performance of the models is reported using root mean squared error (RMSE), median absolute error (MAE), and the Pearson correlation coefficient (r).

### Annotation

OCRs were annotated as promoters if they lay 1000 bp upstream or downstream of the transcription start sites. OCRs were annotated as enhancers if they overlapped regions annotated as enhancers in the PEREGRINE dataset [27]. Notably, we allowed OCRs to be annotated both as promoters and enhancers. In the case of promoters, OCRs were linked to the closest gene, whereas for enhancers, links to genes were sourced from the PEREGRINE dataset.

---

**Supplementary Figure 1.**
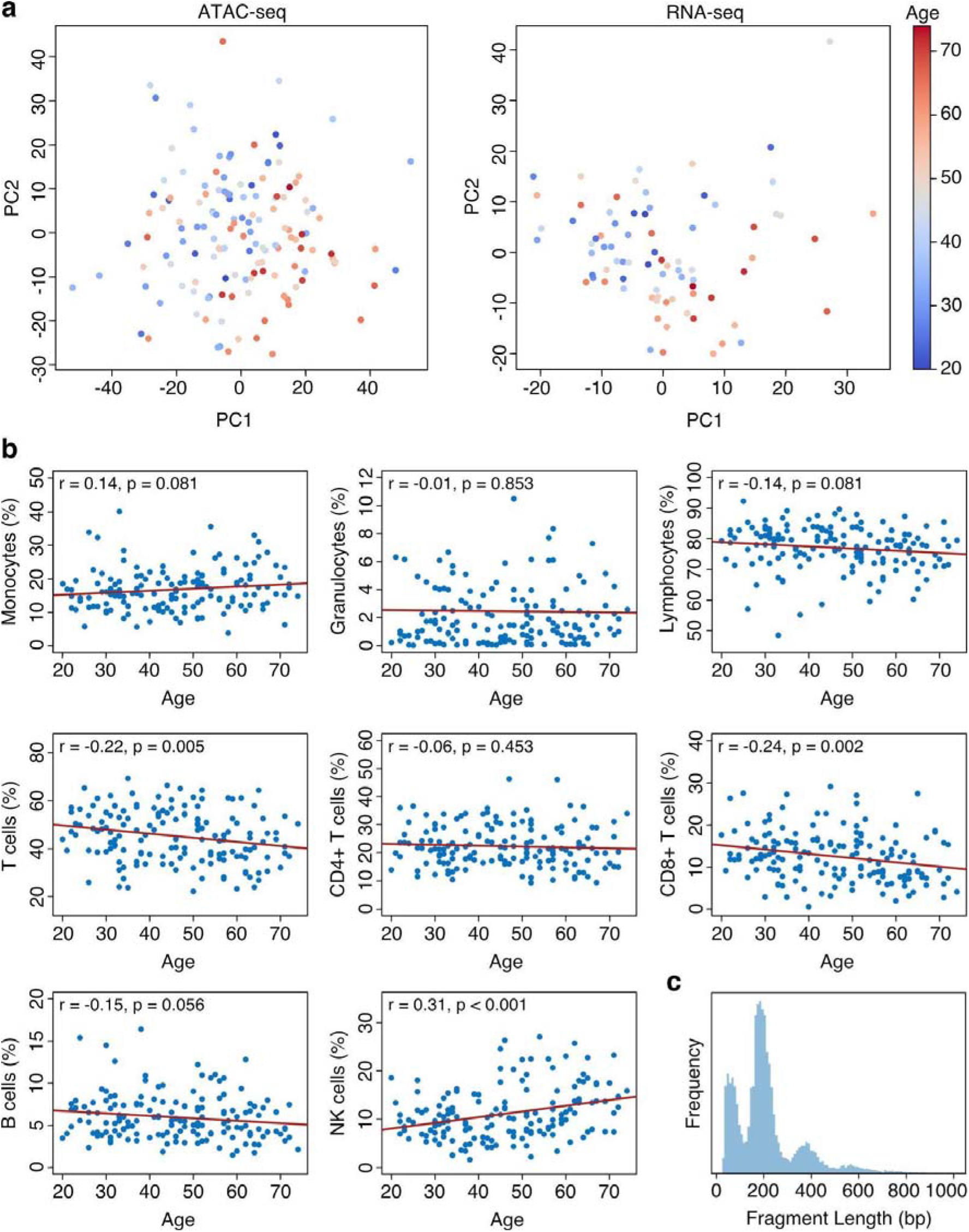
Characterization of ATAC-seq, RNA-seq and flow cytometry data. **a** Principal component analysis (PCA) of the ATAC-seq and RNA-seq data. Principal components 1 and 2 are shown. **b** Changes in immune cell populations with age. **c** Histogram of fragment length distribution for a representative ATAC-seq profile.

**Supplementary Figure 2.**
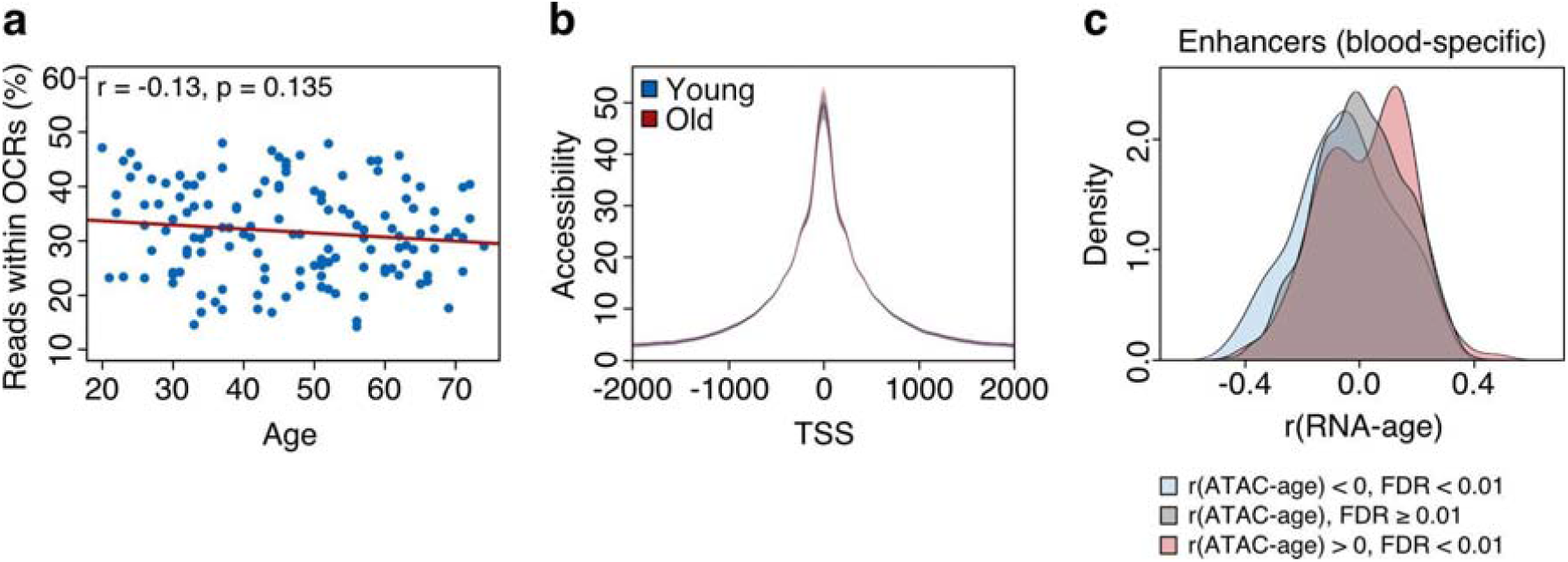
Exploration of global accessibility changes during age. Relationship between accessibility at blood-specific enhancers and transcription. **a** Fraction of reads within peaks plotted against age. **b** Average ATAC-seq coverage profile around TSS for samples of young donors (<35 years, n = 40) and samples of old donors (>55 years, n = 40). **c** Distribution of correlation between transcription levels of genes and age linked to blood-specific enhancers whose accessibility is positively correlated with age (r > 0, FDR < 0.01), negatively correlated with age (r < 0, FDR < 0.01) or not correlated with age (FDR ≥ 0.01).

**Supplementary Figure 3.**
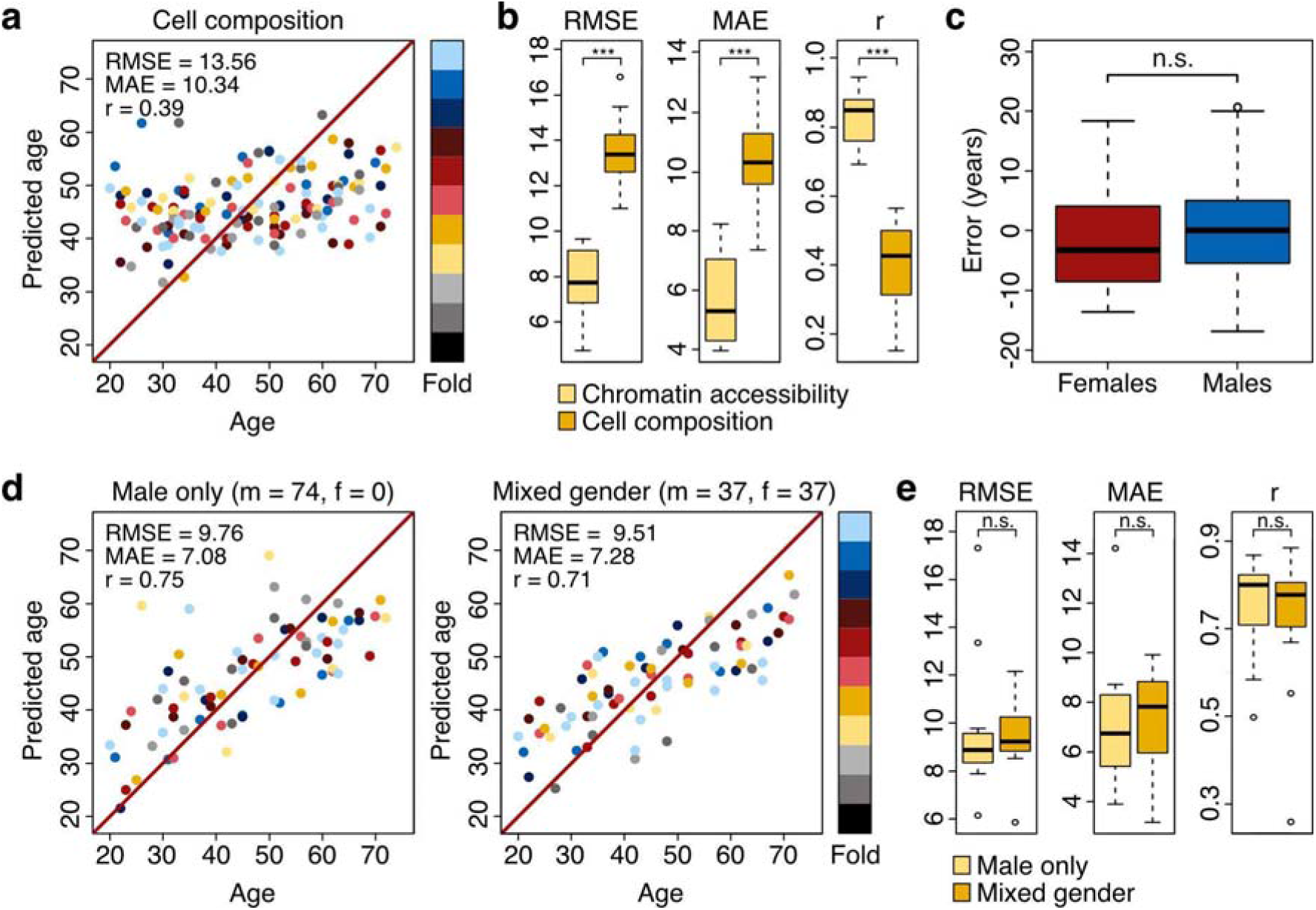
Relevance of cell composition and gender. **a** Development of an aging clock based on cellular composition using nested-cross validation with 11 outer folds (11 different models, each fold in the test set once). Test set predictions from each fold are shown. **b** Performance comparison of chromatin accessibility clock and cell composition clock. **c** Prediction error of chromatin accessibility clock on male and female samples. **d** Comparison of a chromatin accessibility clock trained on male samples only (m = 74, f = 0) to one made with mixed-gender samples (m = 37, f = 37). **e** Performance comparison of male-only and mixed-gender chromatin accessibility clocks.

